# Organelle proteomics reveals novel metabolic vulnerabilities in FLT3-ITD cells

**DOI:** 10.64898/2026.01.19.700272

**Authors:** Valeria Bica, Anna Francesca Pacilè, Martin Boettcher, Valentina Marano, Veronica Marabitti, Francesca Nazio, Mirko Cortese, Thomas Fischer, Livia Perfetto, Dimitrios Mougiakakos, Giorgia Massacci, Francesca Sacco

## Abstract

In acute myeloid leukemia (AML), the insertion site of internal tandem duplications (ITDs) within the *FLT3* gene critically determines the sensitivity to tyrosine kinase inhibitors (TKIs). Despite recent advances, patients harboring ITDs in the tyrosine kinase domain (TKD) still lack effective therapeutic options. To elucidate the molecular basis underlying the differential TKI sensitivity of FLT3-ITD cells, we integrated high-resolution mass spectrometry–based (phospho)proteomics with subcellular fractionation. Our analysis revealed that midostaurin induces the subcellular redistribution of approximately 2500 proteins involved in crucial biological processes, including cell cycle control, autophagy, and metabolism. Functional analyses further demonstrated that the ITD insertion site determines the autophagy response to midostaurin and modulates mitochondrial metabolism, influencing organelle architecture and ATP production, even at steady state. Importantly, by integrating subcellular proteomic dataset with functional metabolic assays, we uncovered a lipid-dependent vulnerability of FLT3-ITD cells: lipid restriction enhances FLT3 trafficking to the plasma membrane, and markedly reduces cell viability, restoring midostaurin sensitivity of resistant FLT3-ITD cells. Together, our findings reveal that the FLT3-ITD insertion site orchestrates a coordinated remodeling of subcellular protein organization, autophagy, and metabolism, and identify lipid-mediated control of FLT3 compartmentalization as a therapeutically actionable mechanism to overcome TKI resistance in FLT3-ITD AML.

## Introduction

Internal tandem duplications (ITDs) in the *FLT3* gene are detected in approximately 25–30% of younger adults with newly diagnosed acute myeloid leukemia (AML) and are associated with an adverse prognosis (1). These mutations most commonly involve exons 14–15, affecting the juxtamembrane domain (JMD) or the first tyrosine kinase domain (TKD1), and resulting in constitutive FLT3 activation through disruption of its autoinhibitory conformation (2). The implementation of the multikinase inhibitor midostaurin to standard induction and consolidation chemotherapy has improved outcomes for patients with *FLT3*-mutated AML (3,4). However, an ever-growing number of clinical and experimental evidence indicates that the prognostic and therapeutic impact of *FLT3*-ITD is influenced by the ITD insertion site (5). While patients harboring JMD insertions (FLT3^ITD-JMD^) benefit from FLT3 inhibition, ITDs within the TKD region (FLT3^ITD-TKD^) are associated with reduced sensitivity to both chemotherapy and FLT3 tyrosine kinase inhibitors (TKIs), as well as lower relapse-free and overall survival (5). Consistent with these observations, experimental models demonstrate that FLT3^ITD-TKD^ mutations confer resistance to apoptosis in response to multiple TKIs, including gilteritinib and quizartinib, and to cytarabine, compared with FLT3^ITD-JMD^ cells (6–8). We recently demonstrated that the impact of ITD insertion site on TKI response is mediated by widespread phosphorylation changes, occurring independently of alterations in mRNA expression or protein abundance (6). Based on these findings, we hypothesized that differential sensitivity of FLT3^ITD-JMD^ and FLT3^ITD-TKD^ cells to midostaurin is driven by phosphorylation-dependent redistribution of the proteome across subcellular compartments. To test this, we performed high-resolution mass spectrometry (MS)–based proteomics combined with subcellular fractionation. This approach identified approximately 1,800 proteins exhibiting genotype-dependent changes in localization following midostaurin treatment. Notably, protein complexes involved in cell-cycle progression, DNA replication, and mitosis translocated from the nucleus to the cytosol exclusively in FLT3^ITD-JMD^ cells, underscoring the profound impact of ITD location on TKI-mediated cell-cycle control. Integration with phosphoproteomic data further revealed ITD-dependent differences in protein phosphorylation and trafficking, highlighting a different regulation of autophagic processes and distinct metabolic shifts in response to midostaurin. Specifically, we found that midostaurin induced autophagy and reduced oxidative phosphorylation and glycolysis in FLT3^ITD-JMD^ but not in FLT3^ITD-TKD^ cells. Furthermore, pharmacological inhibition of key metabolic pathways revealed that midostaurin-treated FLT3^ITD-TKD^ cells retain a partial dependence on lipid metabolism. Notably, lipid deprivation sensitized FLT3^ITD-TKD^ cells to TKI treatment, uncovering a novel metabolic vulnerability in FLT3-ITD AML cells. Overall, our work highlights the importance of integrating proteomic, phosphoproteomic, and metabolic analyses to uncover context-specific vulnerabilities in AML and offers a rationale for developing strategies aimed at overcoming resistance in high-risk FLT3-ITD patients.

## Methods

### Cell culture

Mouse Ba/F3 cells expressing ITD-JMD and ITD-TKD constructs were provided by courtesy of T. Fischer. The cells were cultured in RPMI 1640 medium (Hyclone, Thermo Scientific, Waltham, MA) supplemented with 10% heat-inactivated fetal bovine serum (ECS0090D Euroclone, Italy, MI), 100 U/ml penicillin and 100 mg/ml streptomycin (Gibco 15140122), 1 mM sodium pyruvate (Sigma-Aldrich, St. Louis, Missouri, United States, S8636) and 10 mM 4-(2-hydroxyethyl)-1-piperazineethanesulfonic acid (HEPES) (Sigma H0887). Midostaurin (Selleck chemical, S8064) was used at 100 nM for 24 hours.

Additional methods were included in Supplementary Materials.

## Results

### The experimental strategy

To investigate how the FLT3-ITD insertion site shapes midostaurin-dependent proteome reorganization, we applied a streamlined spatial proteomics workflow (9) combining chemical fractionation with MS-based proteomics (**Fig.1A**). This approach enables high-resolution mapping of protein relocalization across subcellular compartments and provides a quantitative, system-wide view of spatial changes in response to stimuli. Ba/F3 cells expressing FLT3^ITD-JMD^ and FLT3^ITD-TKD^ receptors were treated with 100 nM midostaurin (PKC412) for 24 hours or left untreated. Sequential extraction using six buffers generated six subcellular fractions, each analyzed in biological quadruplicates. Label-free quantification yielded a comprehensive dataset of 8334 proteins, with ∼5000 proteins consistently quantified across fractions (Pearson correlation 0.94–0.99; **Fig.S1A–D**; **Table S1**). As subcellular localization is dynamically regulated by post-translational modifications, we additionally profiled the global phosphoproteome in parallel (**Fig.1A**). This analysis quantified (9)marker proteins displayed precise distribution patterns across fractions as expected for this established protocol (9) : cytosolic markers were enriched in fraction 1; nuclear proteins in fractions 2 and 6; and lysosomal, Golgi, ER, mitochondrial, and endosomal proteins peaked in fractions 3–4 (**Fig.1B**). Consistent with previous reports indicating perinuclear retention of FLT3-ITD mutants (10,11), both FLT3^ITD-JMD^ and FLT3^ITD-TKD^ receptors showed maximal signal in fractions 3–4, corresponding to membrane-bound organelles (**Fig.S2B**). Clustering of protein abundance profiles across fractions revealed five major clusters corresponding to discrete subcellular compartments, with high concordance across genotypes and treatment conditions **(Fig.1C**). Three principal compartment groups were evident: cytosolic and cytoskeletal proteins (fraction 1); membrane-associated proteins, including ER/Golgi and other organelles (fractions 3–4); and nuclear proteins (fractions 2, 5, and 6). Together, these data demonstrate the high reproducibility and resolution of our spatial proteomics pipeline and establish a robust framework to dissect how the FLT3-ITD insertion site governs midostaurin-dependent proteome redistribution and downstream cellular responses, impacting on TKI sensitivity.

**Figure 1.**
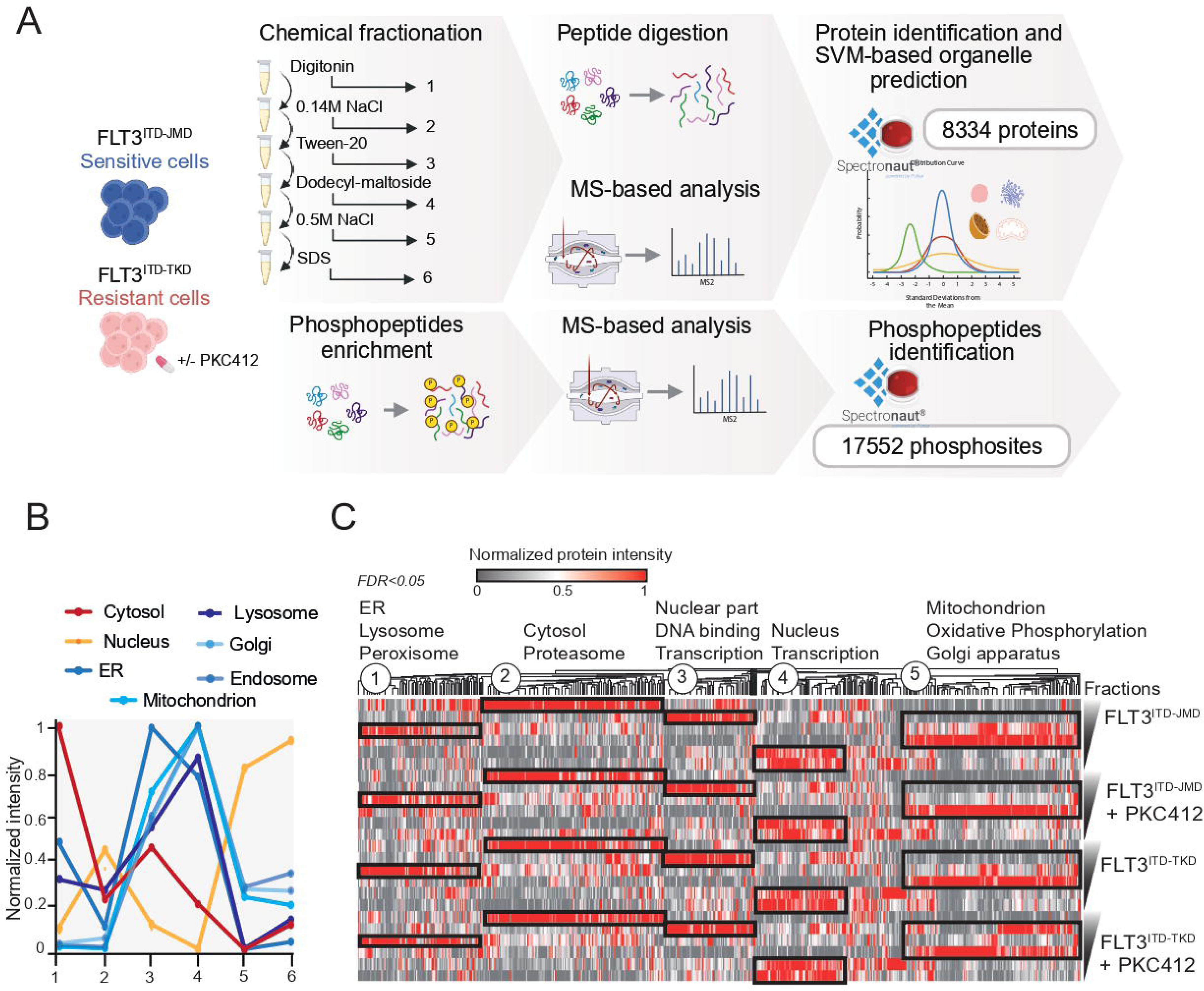
Organelle proteomics generates midostaurin-specific subcellular maps in FLT3-ITD cells. **(A)** Schematic overview of the organelle proteomics pipeline. FLT3^ITD-JMD^ (TKI-sensitive) and FLT3^ITD-TKD^ (TKI-resistant) Ba/F3 cells were left untreated or treated with 100 nM midostaurin for 24 h. Sequential extraction with six buffers generated six subcellular fractions, which were analyzed by MS-based proteomics in biological quadruplicates. Peptides were subjected to global proteomic profiling and phosphopeptide enrichment. **(B)** Intensity profiles of organelle-specific marker proteins across the six fractions. **(C)** Hierarchical clustering of protein profiles (scaled intensities) across fractions for untreated and midostaurin-treated FLT3-ITD cells. Statistically significant enriched gene ontology terms in each cluster are indicated.

### Midostaurin-dependent subcellular redistribution of the global proteome in FLT3-ITD cells

Our spatial proteomic workflow enables accurate quantification of proteins across three major subcellular compartments (cytosol, nucleus, and organelles) in both untreated and midostaurin-treated FLT3-ITD cells (**Fig.1**). To systematically identify proteins translocating between compartments, we applied a classification strategy integrating support vector machine (SVM) learning with correlation profiling (**Fig.2A**). Using compartment-specific marker profiles to train the SVM, we reliably assigned each of the ∼8,000 quantified proteins to a nuclear, cytosolic, or organellar category in every experimental condition (**Fig.2B**; **Table S4**). The high assignment accuracy (>80% of training proteins correctly classified; **Fig.S2C**) confirmed the robustness of our approach.

**Figure 2.**
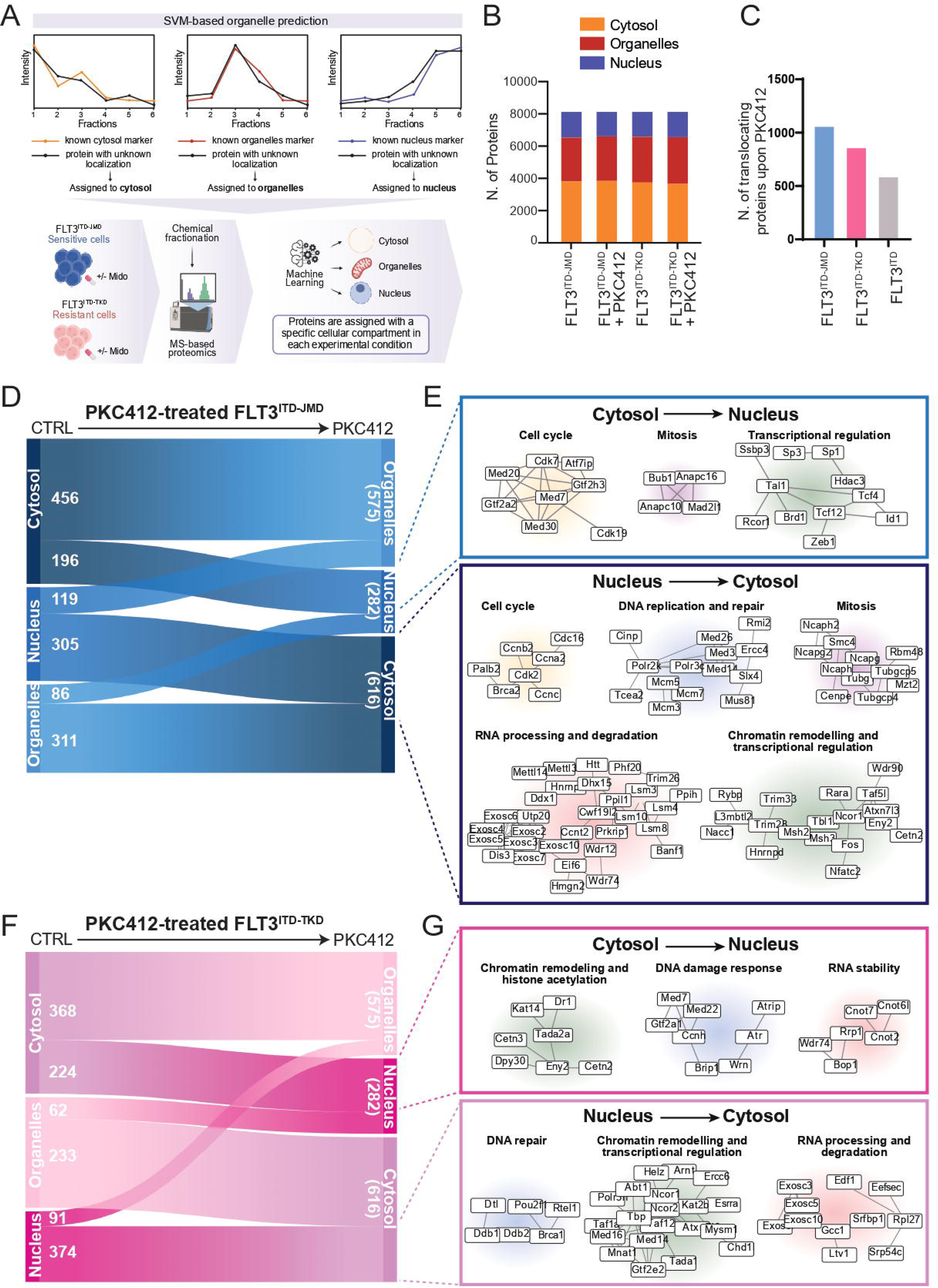
Midostaurin induces genotype-specific subcellular proteome reorganization in FLT3-ITD cells. **(A)** Schematic representation of the bioinformatic strategy used to assign a cell compartment to proteins by using the SVM-algorithm. **(B)** Number of proteins assigned to cytosolic, nuclear, or organellar compartments in untreated and midostaurin treated FLT3-ITD cells. **(C)** Quantification of proteins changing their subcellular localization upon midostaurin treatment. **(D)** Sankey plot of protein translocation in midostaurin-treated FLT3^ITD-JMD^. **(E)** Functional protein complexes redistributed between cytosol and nucleus in midostaurin-treated FLT3^ITD-JMD^. **(F)** Sankey plot of protein translocation in midostaurin-treated FLT3^ITD-TKD^ cells. **(G)** Functional protein complexes redistributed between cytosol and nucleus in midostaurin-treated FLT3^ITD-TKD^ cells.

Across all conditions, 2524 proteins changed their subcellular localization in response to midostaurin. Of these, ∼1065 relocalized specifically in FLT3^ITD-JMD^ cells and 866 in FLT3^ITD-TKD^ cells, while 593 proteins were shared (**Fig.2C**). Notably, only one-third of the shared proteins moved to the same destination compartment, whereas two-thirds redistributed in opposite directions (**Fig.S2D**), underscoring the strong influence of the ITD insertion site on midostaurin-dependent spatial proteome remodeling. We next examined whether specific protein complexes were differentially mobilized between the cytosol and nucleus. Strikingly, FLT3^ITD-JMD^ and FLT3^ITD-TKD^ cells displayed profoundly distinct patterns of complex translocation (**Fig.2D–G**). In midostaurin-sensitive FLT3^ITD-JMD^ cells, treatment induced extensive nuclear export of complexes critical for cell-cycle progression, DNA replication, and mitosis, including CDK–cyclin modules and the MCM helicase complexes (**Fig.2D-E**). Key transcriptional regulators with established roles in AML survival—including NCOR1, RARA, FOS, and TBL1XR1—also redistributed from the nucleus to the cytosol, consistent with disruption of essential survival programs and facilitation of apoptosis. In midostaurin-resistant FLT3 cells, protein-complex translocation was markedly attenuated and qualitatively distinct (**Fig.2F–G**). Relocalized proteins were largely restricted to chromatin-remodeling, histone-regulatory, and RNA-stability pathways. Nuclear import of factors involved in the DNA damage response, including ATR-associated components, suggested activation of protective DNA repair programs. Importantly, major complexes extensively mobilized in FLT3^ITD-JMD^ cells, such as those governing replication licensing, mitotic entry, and cell-cycle progression, remained largely static in FLT3^ITD-TKD^cells.

### Genotype-specific autophagy response in midostaurin-treated FLT3-ITD cells

To investigate whether midostaurin-dependent translocating proteins in FLT3-ITD cells were involved in additional specific cellular processes, we performed a Gene Ontology enrichment analysis. Consistently with our data on protein complexes, proteins involved in cell cycle progression, DNA replication and repair were mostly translocated between cytoplasm and nucleus in FLT3^ITD-JMD^ cells and, to a lesser extent, in FLT3^ITD-TKD^ cells. Furthermore, our analysis showed a relocalization of autophagy-related proteins in both FLT3^ITD-JMD^ and FLT3^ITD-TKD^ cells following midostaurin treatment (**Fig.3A–B**). A more detailed investigation revealed pronounced differences in the subcellular distribution of these proteins between TKI-sensitive and resistant FLT3-ITD cells. In FLT3^ITD-JMD^ cells, midostaurin treatment led to the recruitment of the TFEB–FNIP–FLCN complex and the adaptor protein p62 (SQSTM1) to an organellar compartment, most likely corresponding to lysosomes (14,15) (**Fig.3C**). This observation suggests that midostaurin treatment likely promotes autophagy in FLT3^ITD-JMD^ cells. In contrast, in FLT3^ITD-TKD^ cells, the TFEB–FNIP–FLCN complex and p62 lost their association with these organelles after treatment and instead became diffusely distributed throughout the cytoplasm (**Fig.3C**). Moreover, by integrating the subcellular proteomic data with the phosphoproteomic profiles (**Fig.S2E**), we found that the autophagy-related proteins displaying differential subcellular localization in FLT3^ITD-JMD^ versus FLT3^ITD-TKD^ cells, also showed opposite phosphorylation patterns after midostaurin treatment (**Fig.3D**). Except for TFEB (S466) and FLCN (S62), the regulatory roles of the other phosphosites are currently undefined. Nevertheless, our analysis suggests that these phosphosites may participate in controlling protein intracellular localization. These observations led us to hypothesize that midostaurin treatment differentially affects autophagy in FLT3-ITD sensitive and resistant cells. To this aim, we measured autophagy flux in both sensitive and resistant FLT3-ITD cells treated with midostaurin, either alone or in combination with Bafilomycin A1, an inhibitor of lysosomal V-ATPase. As shown by the accumulation of LC3-II, midostaurin treatment induced autophagy only in FLT3^ITD-JMD^ cells (**Fig.3E**). We next investigated whether FLT3^ITD-TKD^ cells are intrinsically defective in autophagy. Thus, FLT3^ITD-JMD^ and FLT3^ITD-TKD^ cells were starved for 3 hours in the presence or absence of Bafilomycin A1, and autophagy flux was monitored by measuring LC3 lipidation. Both FLT3-ITD cell types showed comparable induction of autophagy upon starvation, indicating that midostaurin specifically affects autophagy in a FLT3 genotype-specific manner (**Fig.S2F**). As autophagy impairment has been linked to cancer drug resistance (16), we investigated whether autophagy induction could increase the TKI sensitivity of FLT3^ITD-TKD^ cells. To this aim, FLT3-ITD cells were exposed to midostaurin for 24 hours, nutrient deprivation for 3 hours, or a combination of both treatments. Strikingly, starvation restored the sensitivity of FLT3^ITD-TKD^ cells to midostaurin, suggesting that impaired autophagy may contribute to their resistance phenotype (**Fig.3F**). These findings demonstrate that the ITD insertion site influences the capacity of AML cells to activate autophagy in response to midostaurin. Moreover, our data show that short-term nutrient deprivation induces cell death in both FLT3-ITD cell types, highlighting a previously unrecognized potential metabolic vulnerability.

**Figure 3.**
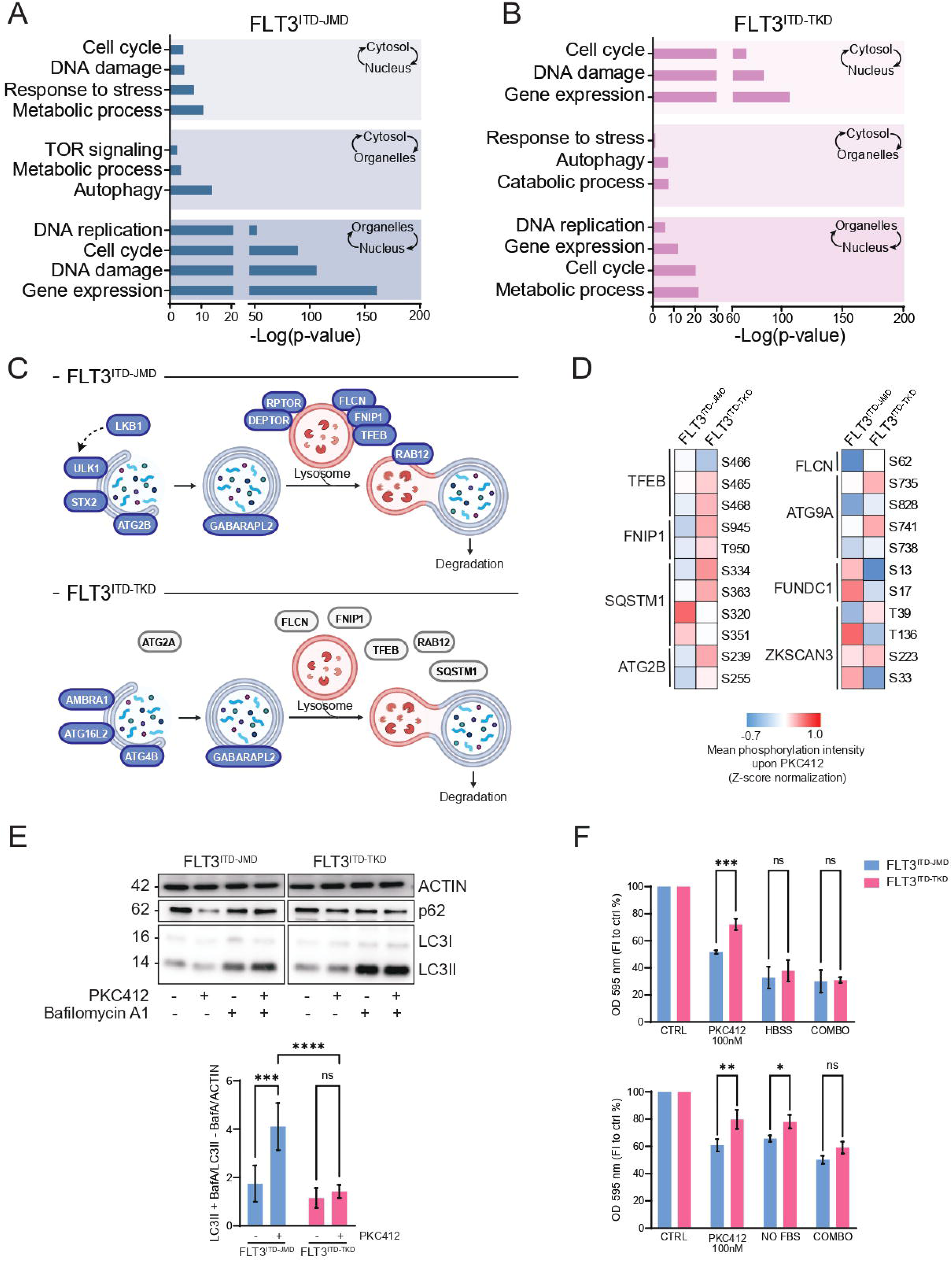
Midostaurin-induced mitochondrial remodelling and spatial changes. (A-B) Bar plots reporting the -log2 (p-value) of the GO-Term enrichment analysis of translocating proteins upon midostaurin treatment in sensitive and resistant FLT3-ITD cells. **(C)** Schematic representation of the spatial remodelling of key autophagic proteins upon midostaurin treatment in sensitive and resistant FLT3-ITD cells. **(D)** Heatmap reporting the intensity of the phosphorylation level of translocating autophagic proteins after midostaurin treatment in FLT3^ITD-JMD^ and FLT3^ITD-TKD^. **(E)** Western blotting of LC3-II and p62 protein levels in FLT3^ITD-JMD^ and FLT3^ITD-TKD^. Cells were treated with midostaurin with or without bafilomycin A1 for 24 hours. **(F)** Bar plot reporting the percentage of viable cells measured by MTT assay. FLT3^ITD-JMD^ and FLT3^ITD-TKD^ were treated with 100nm of midostaurin treatment for 24h either alone or combined with 3 hours of starvation.

### Midostaurin induces ITD site–dependent metabolic reprogramming

Our results showed that the ITD insertion site determines the autophagy response to midostaurin and uncover a shared vulnerability of both FLT3-ITD cell types to short-term nutrient deprivation. Building on these observations, we next investigated how ITD location influences the metabolic response to midostaurin. To this end, FLT3^ITD-JMD^ and FLT3^ITD-TKD^ cells were treated with midostaurin for 24h or left untreated and their glycolytic and oxidative phosphorylation activities were assessed using Seahorse extracellular flux analysis. In control condition, FLT3^ITD-TKD^ cells displayed a slightly higher glycolysis and glycolytic capacity than FLT3^ITD-JMD^ cells, as indicated by elevated extracellular acidification rate (ECAR) (**Fig.4A**). This metabolic profile was further supported by significantly elevated basal respiration and ATP production in FLT3^ITD-JMD^ cells compared to FLT3^ITD-TKD^ cells (**Fig.4B**). Consistently, transmission electron microscopy (TEM) images revealed that FLT3^ITD-JMD^ cells appear to have a higher number of cristae and an increased cristae-to-mitochondrial ratio compared to FLT3^ITD-TKD^ cells, a morphology in line with their higher basal respiration and ATP production (**Fig.4C-D**). Following midostaurin (PKC412) treatment, FLT3^ITD-JMD^ cells displayed a strong reduction in glycolysis, glycolytic capacity, basal respiration and ATP production (**Fig.4A-B**). In contrast, FLT3^ITD-TKD^ cells did not exhibit significant metabolic changes upon midostaurin exposure. Consistently, phosphorylation levels of PKM2 Y105 decreased upon midostaurin treatment only in FLT3^ITD-JMD^ cells, but not in FLT3^ITD-TKD^ cells (**Fig.S2G**). Together, these results demonstrate that the ITD insertion site influences key metabolic processes already under steady-state conditions and highlights the metabolically TKI-resistant phenotype of FLT3^ITD-TKD^ cells.

**Figure 4.**
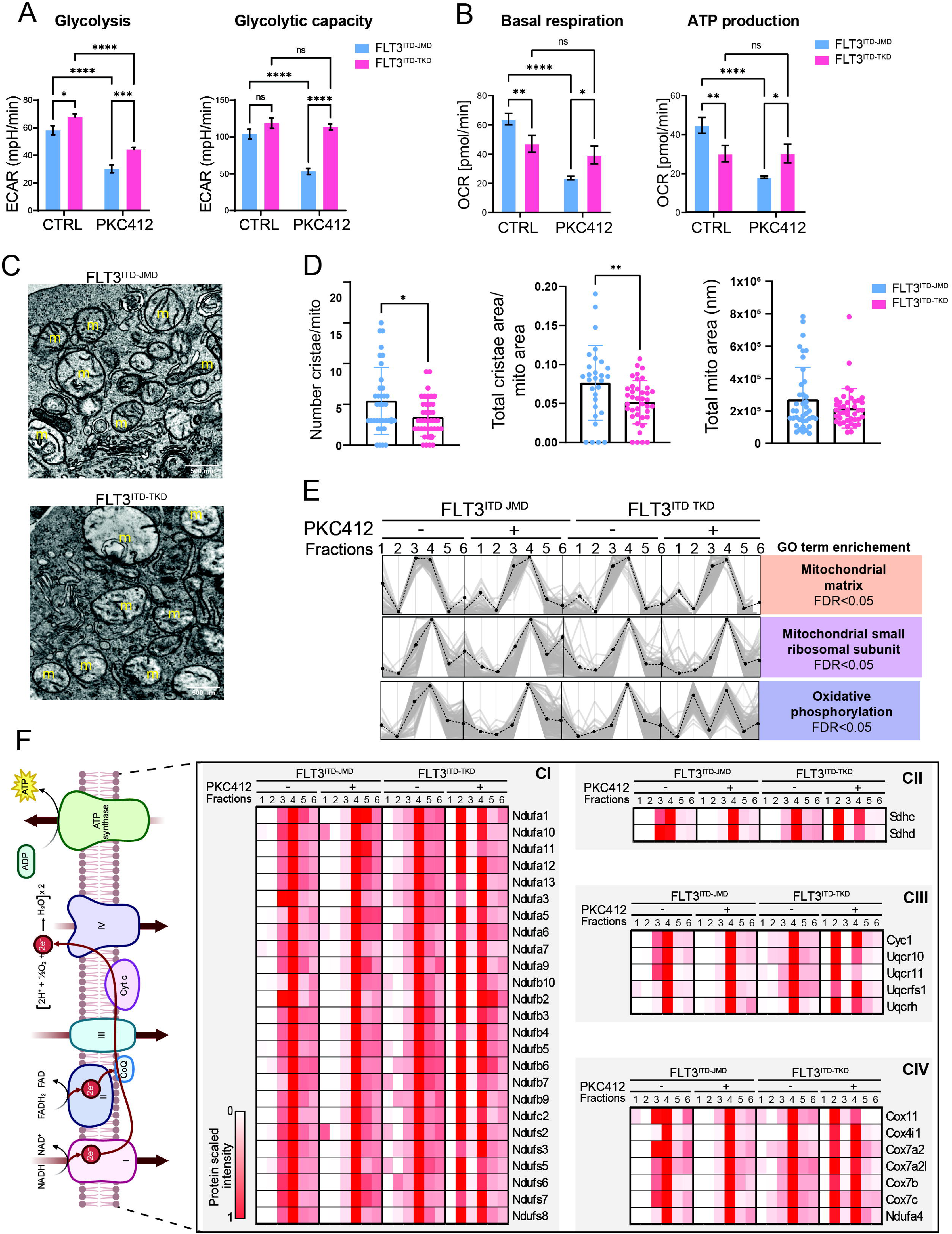
Midostaurin induces ITD site–dependent metabolic reprogramming. **(A)** Bar plot reporting the ECAR based measurements of glycolysis and glycolytic capacity in FLT3^ITD-JMD^ and FLT3^ITD-TKD^. Cells were treated with 100nm midostaurin for 24 hours and subjected to a glycolysis stress test. (**B**) OCR-derived assessments of basal respiration and ATP production in FLT3^ITD-JMD^ and FLT3^ITD-TKD^. Cells were treated with 100nm midostaurin for 24 hours and subjected to a mitochondrial stress test. **(C)** Representative electron microscopy images of steady-state FLT3^ITD-JMD^ and FLT3^ITD-TKD^ cells (m=mitochondria). **(D)** Bar plots reporting the number of cristae, the total cristae area and total mitochondria area in steady-state FLT3^ITD-JMD^ and FLT3^ITD-TKD^ cells. **(E)** Intensity profiles of the 168 proteins annotated in the MitoCarta database as mitochondrial proteins across the six fractions in each experimental condition. Colored boxes indicate the enriched GO-Term biological processes and cell compartments. **(F)** Heatmaps reporting the protein levels (scaled intensities) of the OXPHOS related proteins across the 6 fractions in each experimental condition.

Next, we leveraged our subcellular proteomics dataset to examine how mitochondrial proteins distribute across fractions. We focused on the 168 proteins annotated as “mitochondrial” in the MitoCarta database (17) and analyzed their abundance profiles across the six fractions. Unsupervised clustering identified three major groups enriched for mitochondrial matrix proteins, mitochondrial small ribosomal subunits, and oxidative phosphorylation (OXPHOS) complexes (**Fig.4E**). Mitochondrial matrix proteins and small ribosomal subunit proteins displayed highly similar distribution profiles in untreated and midostaurin-treated FLT3-ITD cells, indicating that these mitochondrial components are not substantially affected neither by TKI exposure nor by the ITD insertion site (**Fig.4E)**. In contrast, OXPHOS enzymes showed a markedly different behavior. Strikingly, only in midostaurin-treated FLT3^ITD-TKD^ cells the OXPHOS complex I–IV subunits shift their distribution, exhibiting a prominent enrichment in fraction 2 (nuclear compartment) (**Fig.4F**). This redistribution was not observed in FLT3^ITD-JMD^ cells, highlighting a ITD-specific alteration of the mitochondrial respiratory machinery upon midostaurin treatment.

### Lipid–dependent mitochondrial activity in FLT3^ITD-TKD^ cells

Our results revealed that FLT3^ITD-TKD^cells are metabolically unresponsive to TKI treatment (i) (**Fig.4A-B**), exhibit an altered distribution of their OXPHOS complexes (ii) (**Fig.4E-F**), and can be re-sensitized to TKI by short-term nutrient deprivation (iii) (**Fig.3F**). These findings prompted us to investigate which metabolic substrates sustain mitochondrial respiration in FLT3^ITD-TKD^ cells under midostaurin treatment. Subcellular proteomics showed that both FLT3^ITD-JMD^ and FLT3^ITD-TKD^ cells exhibit a strong enrichment in enzymes involved in lipid metabolism translocating from cytosol to organelles (**Fig.5A**), suggesting alternative substrate utilization.

**Figure 5.**
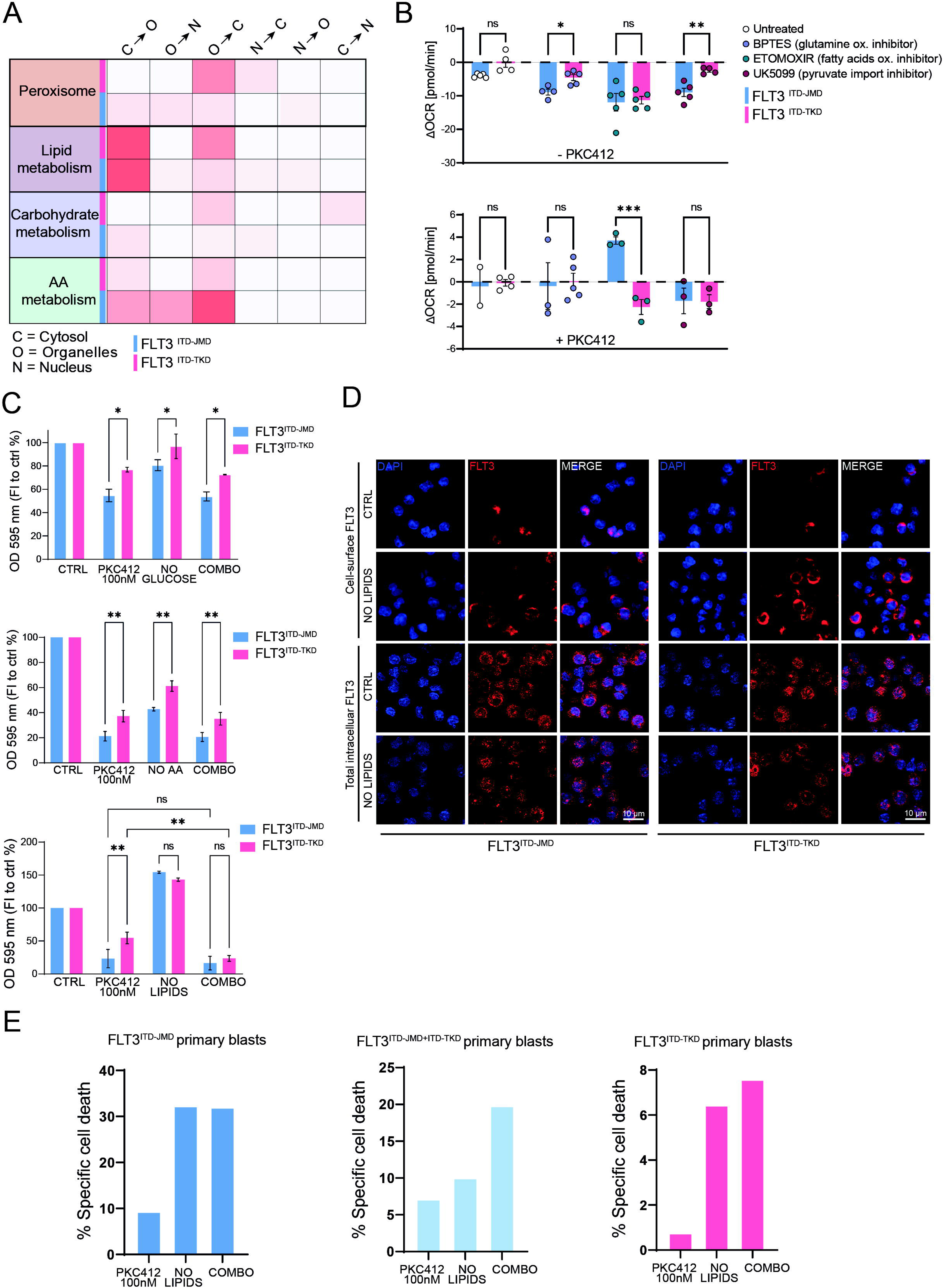
Lipid availability modulates FLT3-ITD localization and TKI response. **(A)** Heatmap reporting the number of translocating proteins involved in carbohydrate, amino acids, peroxisome and lipid metabolism in FLT3^ITD-JMD^ and FLT3^ITD-TKD^. **(B)** Bar plot reporting the ΔOCR of basal respiration after the inhibition of glucose usage (UK5099), glutamine usage (BPTES) and lipid usage (ETOMOXIR) for OXPHOS in FLT3^ITD-JMD^ and FLT3^ITD-TKD^ in absence and presence of 100nM midostaurin treatment for 24h. **(C)** Bar plot reporting the percentage of viable cells measured by MTT assay. FLT3^ITD-JMD^ and FLT3^ITD-TKD^ were treated with 100nm of midostaurin treatment for 24h either alone or combined with glucose, amino acid and lipid deprivation. **(D)** Representative immunofluorescence images of FLT3 localization before and after lipid deprivation in both permeabilized and non-permealized FLT3^ITD-JMD^ and FLT3^ITD-TKD^. **(E)** Barplots showing the percentage of treatment-induced apoptosis (100 * (dead cells after treatment – death cells in control) / viable cells in control) in patient-derived blasts carrying FLT3-ITD in i) the JM domain, ii) both the JM and TK domain and iii) the TK domain upon the indicated treatments. Percentage of apoptotic cells was assessed by Annexin-V labeling.

To investigate how FLT3-ITD cells fuel mitochondrial respiration, we examined changes in oxygen consumption rate (ΔOCR) following inhibition of glutamine oxidation (BPTES), fatty-acid β-oxidation (Etomoxir), and pyruvate import (UK5099) in FLT3^ITD-JMD^ and FLT3^ITD-TKD^ cells, with or without midostaurin treatment (**Fig.5B**). As previously shown (**Fig.4A-B**), midostaurin markedly suppresses ATP production in FLT3^ITD-JMD^ cells, whereas FLT3^ITD-TKD^ cells remain metabolically insensitive. Inhibition of mitochondrial fuel pathways, in absence of midostaurin, revealed distinct metabolic features of FLT3^ITD-TKD^cells. Blockade of glutamine oxidation and pyruvate import resulted in a stronger ΔOCR reduction in FLT3^ITD-JMD^ compared with FLT3^ITD-TKD^ cells, suggesting a higher dependence on glucose and glutamine metabolism in the JMD context. Fatty-acid oxidation inhibition causes a clear decrease in ΔOCR in both genotypes (**Fig.5B**, upper panel). Conversely, in PKC412 treated cells, etomoxir produces a divergent response: ΔOCR significantly decreases in FLT3^ITD-TKD^ cells, while FLT3^ITD-JMD^ cells display a compensatory increase in respiration (**Fig.5B**, lower panel), demonstrating that FLT3^ITD-TKD^ cells remain critically dependent on fatty-acid oxidation to sustain mitochondrial activity when FLT3 signaling is inhibited.

### Lipid availability modulates FLT3-ITD localization and TKI response

To further explore whether altered fatty-acid metabolism contributes to the differential TKI sensitivity of FLT3-ITD variants, we assessed cell viability under nutrient-restricting conditions, by the removal of glucose, amino acids, or lipids, alone or in combination with midostaurin. Specifically, FLT3^ITD-JMD^and FLT3^ITD-TKD^cells were incubated with or without midostaurin in a medium deprived of glucose, amino acids or lipids for 24h. As shown in **Fig.5C**, FLT3^ITD-JMD^ cells displayed a substantial decrease in viability upon midostaurin treatment, and this effect was not impacted when glucose or amino acids were removed. In contrast, FLT3^ITD-TKD^ cells preserved their intrinsic resistance to TKI treatment and remained largely insensitive to midostaurin under glucose or amino-acid deprivation. However, lipid deprivation produced a different pattern: although removal of lipids alone increased cell viability in both genotypes, its combination with midostaurin specifically abolished midostaurin resistance in FLT3^ITD-TKD^ cells, resulting in a pronounced viability loss (**Fig.5C)**. These findings are consistent with our metabolic analyses showing that, upon FLT3 inhibition, FLT3^ITD-TKD^cells heavily rely on fatty-acid oxidation to sustain residual mitochondrial respiration (**Fig.5B**). Accordingly, restricting lipid availability removes essential metabolic support and selectively restores TKI sensitivity in FLT3^ITD-TKD^cells. Together, these findings indicate that fatty-acid metabolism represents a crucial metabolic axis in FLT3-ITD cells, and that, although FLT3^ITD-TKD^ cells are generally metabolically resistant to TKI treatment, their survival becomes markedly compromised when lipid availability is restricted. This prompted us to investigate more closely how lipid deprivation affects FLT3-ITD biology.

FLT3-ITD receptors are known to undergo S-palmitoylation by the ER-resident palmitoyl acyltransferase ZDHHC6 (18). Unlike wild-type FLT3, which traffics efficiently to the plasma membrane, FLT3-ITD accumulates predominantly in the ER (19). We therefore asked whether lipid deprivation could influence FLT3-ITD cell viability by altering palmitoylation-dependent trafficking and thereby modifying FLT3 subcellular localization.

To test this, we examined FLT3 localization by immunofluorescence in FLT3^ITD-JMD^and FLT3^ITD-TKD^ cells cultured in control (CTRL) or lipid-depleted serum (LD-FBS). Cells were stained either without permeabilization, to assess cell-surface FLT3, or with permeabilization, to visualize total intracellular FLT3 (**Fig.5D**). Under control conditions, both FLT3^ITD-JMD^ and FLT3^ITD-TKD^ cells displayed minimal FLT3 signal at the plasma membrane, consistent with the well-described intracellular retention of FLT3-ITD. Strikingly, lipid deprivation caused a robust increase in cell-surface FLT3 in both genotypes, with a prominent ring-like staining pattern more pronounced in FLT3^ITD-TKD^cells but clearly visible in FLT3^ITD-JMD^cells as well. Because cell-surface FLT3 is known to have stronger oncogenic potential than its ER-retained counterpart (19), the increased membrane localization observed under lipid-deprived conditions is consistent with the enhanced viability seen in the absence of midostaurin. In contrast, when lipid deprivation was combined with midostaurin treatment, FLT3-ITD cells exhibited a marked reduction in viability (**Fig.5C**). This finding suggests that lipid-mediated redistribution of FLT3^ITD-TKD^ to the plasma membrane increases receptor accessibility to midostaurin, thereby restoring TKI sensitivity. Overall, these data suggest that lipid-dependent FLT3-ITD compartmentalization regulates membrane exposure and TKI response, restoring sensitivity in FLT3-ITD-TKD cells and affecting midostaurin-sensitive populations. Importantly, although beyond the scope of the present study, we validated this lipid-dependent modulation of TKI response ex vivo. To this end, we used primary AML blasts carrying different FLT3-ITD mutations: FLT3^ITD-TKD^ blasts (carrying the ITD in the TK domain), FLT3 ^ITD-JMD+ITD-TKD^ blasts (carrying the ITD in both the JM and TK domain), and FLT3^ITD-JMD^ blasts (carrying the ITD in the JM domain). Our experiments showed that lipid deprivation in combination with midostaurin led to a marked increase in cell death in all the FLT3-ITD samples, confirming the translational relevance of lipid dependency in primary leukemic cells (**Fig.5E**). However, it is important to highlight that lipid deprivation alone exerts opposite effects between Ba/F3 cells and primary AML blasts. This discrepancy likely reflects intrinsic differences between the models. Ba/F3 cells overexpress ectopic FLT3-ITD, which may alter signaling and metabolic dependencies and mask metabolic stress responses. In contrast, primary AML blasts represent a more physiologically relevant context, and the endogenous FLT3-ITD expression may confer sensitivity to lipid availability. Collectively, our findings indicate that fatty-acid metabolism represents a critical metabolic axis in FLT3-ITD–driven leukemia and that lipid availability broadly influences TKI responsiveness across FLT3-ITD subtypes, uncovering a shared, lipid-dependent vulnerability that may be therapeutically exploitable (**Fig.6**).

**Figure 6.**
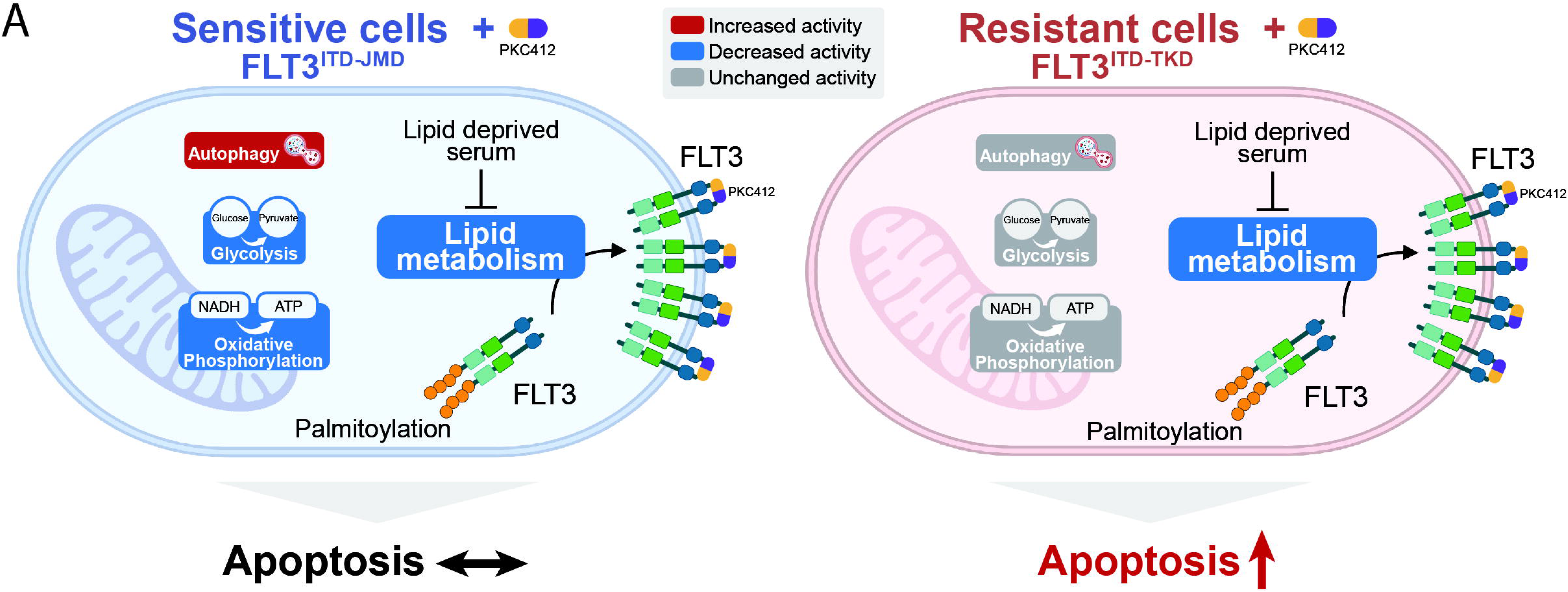
Graphic model representing the metabolic adaptation and vulnerability of FLT3-ITD cells in response to PKC412 treatment.

## Discussion

FLT3-ITD mutations are among the most important genetic determinants of prognosis and therapeutic response in AML (20). Midostaurin in combination with chemotherapy has improved outcomes for FLT3-mutated patients, however a significant subset, especially those harboring ITDs within the TKD region, continues to exhibit primary resistance and poor survival (5).

Our study revealed that the FLT3-ITD insertion site affects how leukemic cells respond to kinase inhibition, by coordinating changes in protein localization, phosphorylation, autophagy, and metabolic plasticity. By using high-resolution subcellular proteomics, we show that midostaurin induces a profound reorganization of the proteome, affecting the localization of more than 2,500 proteins. The functional consequences of such huge proteome remodelling are strongly dictated by the ITD insertion site. In FLT3^ITD-JMD^ cells, proteins critical for DNA replication, cell-cycle progression and mitosis (such as CDK–cyclin complexes, the MCM helicase, and transcriptional regulators) undergo extensive nuclear export, contributing to cell cycle arrest and apoptosis (21). In contrast, FLT3^ITD-TKD^ cells exhibit a different pattern of protein mobilization, including the recruitment of chromatin-modifying and DNA repair proteins to the nucleus, suggesting activation of protective stress-repair response.

One of the most striking findings of our study is that autophagy response to midostaurin is ITD-dependent. Indeed, midostaurin triggers robust autophagy activation only in FLT3^ITD-JMD^ cells, as observed by organellar recruitment of TFEB, FNIP/FLCN and p62, together with increased LC3 lipidation. This pattern is consistent with the well-established mechanism whereby the FNIP/FLCN complex is dynamically recruited to lysosomal membranes, regulating TFEB activation in response to metabolic or lysosomal stress (14). These findings are consistent with recent evidence linking defective autophagy signaling to drug resistance in multiple cancer types (22).

Moreover, our study indicates that ITD location remodels cellular metabolism already under steady-state conditions. Transmission electron microscopy (TEM) combined with metabolic assays revealed that FLT3^ITD-JMD^ cells harbor a greater number of mitochondria that are larger and richer in cristae compared with FLT3^ITD-TKD^ cells, consistent with their higher basal respiration and ATP production (23,24). The molecular mechanisms driving this ITD-dependent mitochondrial and metabolic remodeling remain unknown and require further investigation. In line with their differential sensitivity to TKI treatment, FLT3^ITD-JMD^cells undergo a marked reduction in both glycolysis and oxidative phosphorylation upon midostaurin exposure, coupled with mitochondrial condensation and a consequent drop in ATP levels. By contrast, FLT3^ITD-TKD^ cells maintain stable glycolytic and respiratory activity, displaying a metabolically resilient phenotype.

Integrating our subcellular proteomic dataset with metabolic analyses, we identified lipid metabolism as a potential critical determinant of TKI resistance. Our data show that lipid deprivation, but not amino acid or glucose deprivation, sensitizes FLT3^ITD-TKD^ cells to midostaurin treatment. We hypothesize that this lipid-dependent modulation of TKI sensitivity reflects a direct effect on FLT3. FLT3-ITD palmitoylation by the ER-resident acyltransferase ZDHHC6 promotes receptor retention within the ER (18). Our working model proposes that lipid deprivation reduces palmitoylation efficiency, thereby increasing FLT3 surface exposure. As cell-surface FLT3 possesses higher oncogenic activity and enhances proliferation, its redistribution under lipid-depleted conditions is consistent with the increased viability observed in the absence of midostaurin in cell line models. However, increased membrane localization also renders the receptor more accessible to TKIs, resulting in enhanced cell death in resistant FLT3^ITD-TKD^ cells. Importantly, we validated the lipid vulnerability of FLT3-ITD cells in ex vivo experiments using primary AML blasts. Importantly, ex vivo experiments using primary AML blasts revealed that lipid deprivation alone is sufficient to induce cell death, highlighting an intrinsic lipid dependency of FLT3-ITD–positive blasts. However, given the small number of patient samples included, our findings will require additional confirmation, which lies beyond the scope of the present study. Lipid deprivation has previously been implicated in FLT3-ITD–driven leukemia (25–27). This resulted in clinical trials assessing the potential benefit of introducing low-fat diet regimen or statins to chemotherapy treatment (28). Accordingly, our findings demonstrated that lipid-dependent compartmentalization of FLT3-ITD governs therapeutic vulnerability. However, our data also suggest that lipid deprivation may amplify additional metabolic and signaling vulnerabilities beyond the alteration of receptor localization. In particular, restriction of lipid availability could impair mitochondrial membrane composition and electron transport chain efficiency or affect post-translational lipid modifications of mitochondrial proteins, thereby perturbing pro-survival signaling networks (29–31). Moreover, lipid deprivation can modify membrane composition, affecting receptor organization and signaling and thereby increasing sensitivity to TKI treatment (30,31). Dissecting these interconnected lipid-dependent mechanisms will be essential to fully understand how lipid restriction synergizes with TKIs to overcome resistance. Collectively, our results provide mechanistic insight into how the structural context of the FLT3-ITD mutation influences the spatial proteome landscape and the cellular response to kinase inhibition. The coupling of spatial proteomics with phosphoproteomics offers a powerful framework for dissecting subcellular drug effects and identifying compartment-specific resistance mechanisms in oncogenic signaling.

## Data availability

The mass spectrometry-based proteomics data have been deposited at the ProteomeXchange Consortium, via the PRIDE partner repository.

## Acknowledgments

This research was funded by the Italian Association for Cancer Research (AIRC) with a grant to L.P. (MFAG Grant n. 28858) and a grant to F.S. (Start-Up Grant n. 21815) and by MUR PRIN 2022 (n. E53D23004850006). G.M. is supported by MUR PRIN 2022 (n. E53D23004850006). L.P. and F.S. are supported by a joint PRIN 2022 PNRR grant (n. P2022JRETW), funded by the European Union – NextGenerationEU, and by a SEED Sapienza Grant.

## Author contributions

V.B. and A.F.P. were responsible for the conceptualization of the study together with G.M., L.P. and F.S. The methodology was designed by V.B., A.F.P. and G.M., with contributions from S.G., M.B., L.P., F.N., V.M., M.C. and F.S. Formal analysis was carried out by V.B., A.F.P. and G.M., with the involvement of S.G., M.B., F.N., V.M. and M.C. The investigation was conducted by V.B. and A.F.P., with contributions from T.F., M.B. and D.M. V.B., A.F.P., G.M. and F.S. prepared the original draft of the manuscript. All authors contributed to reviewing and editing the manuscript. Supervision was provided by G.M. and F.S., while F.S. was responsible for funding acquisition. All authors have read and agreed to the published version of the manuscript.

## Competing Interests

The authors declare no competing interests.

---

**Figure S1. (A-D)** Heatmaps reporting the Pearson Correlation coefficient of the proteomic analysis of the subcellular fractionation of control FLT3^ITD-JMD^(A), midostaurin-treated FLT3^ITD-JMD^ (B), FLT3^ITD-TKD^ (C) and midostaurin-treated FLT3^ITD-TKD^ (D). **(E-F)** Bar plots reporting the number of peptides identified in each biological condition and subcellular fraction in FLT3^ITD-JMD^(E) and FLT3^ITD-TKD^(F). **(G)** Heatmap reporting the Pearson Correlation coefficient of the phosphoproteomic analysis of control and midostaurin-treated FLT3^ITD-JMD^ and FLT3^ITD-TKD^.

**Figure S2. (A)** Hierarchical clustering of subcellular fractionation data. **(B)** Intensity profiles of FLT3 protein in the six fractions (scaled intensities). **(C)** Bar plot reporting the percentage of the prediction efficiency of the SVM-organelle assignment. **(D)** Upset plot showing the number of translocating proteins that share the same subcellular location after midostaurin treatment in FLT3^ITD-JMD^ versus FLT3^ITD-TKD^ cells, or unique to each condition. **(E)** Venn diagrams reporting the percentage of translocating proteins whose phosphorylation is significantly modulated by midostaurin treatment or not. **(F)** Representative western blot and relative quantification of LC3-II protein levels after induction of autophagy by starvation in FLT3^ITD-JMD^ and FLT3^ITD-TKD^either alone or combined with midostaurin treatment. (**G**) Representative western blot reporting the phosphorylation level of PKM2 at Y105 in FLT3^ITD-JMD^ and FLT3^ITD-TKD^ after 24 hours of 100nM midostaurin treatment.

## Notes

### Competing Interest Statement

The authors have declared no competing interest.

