## Supplementary figures for "Organelle proteomics reveals novel metabolic vulnerabilities in FLT3-ITD cells"

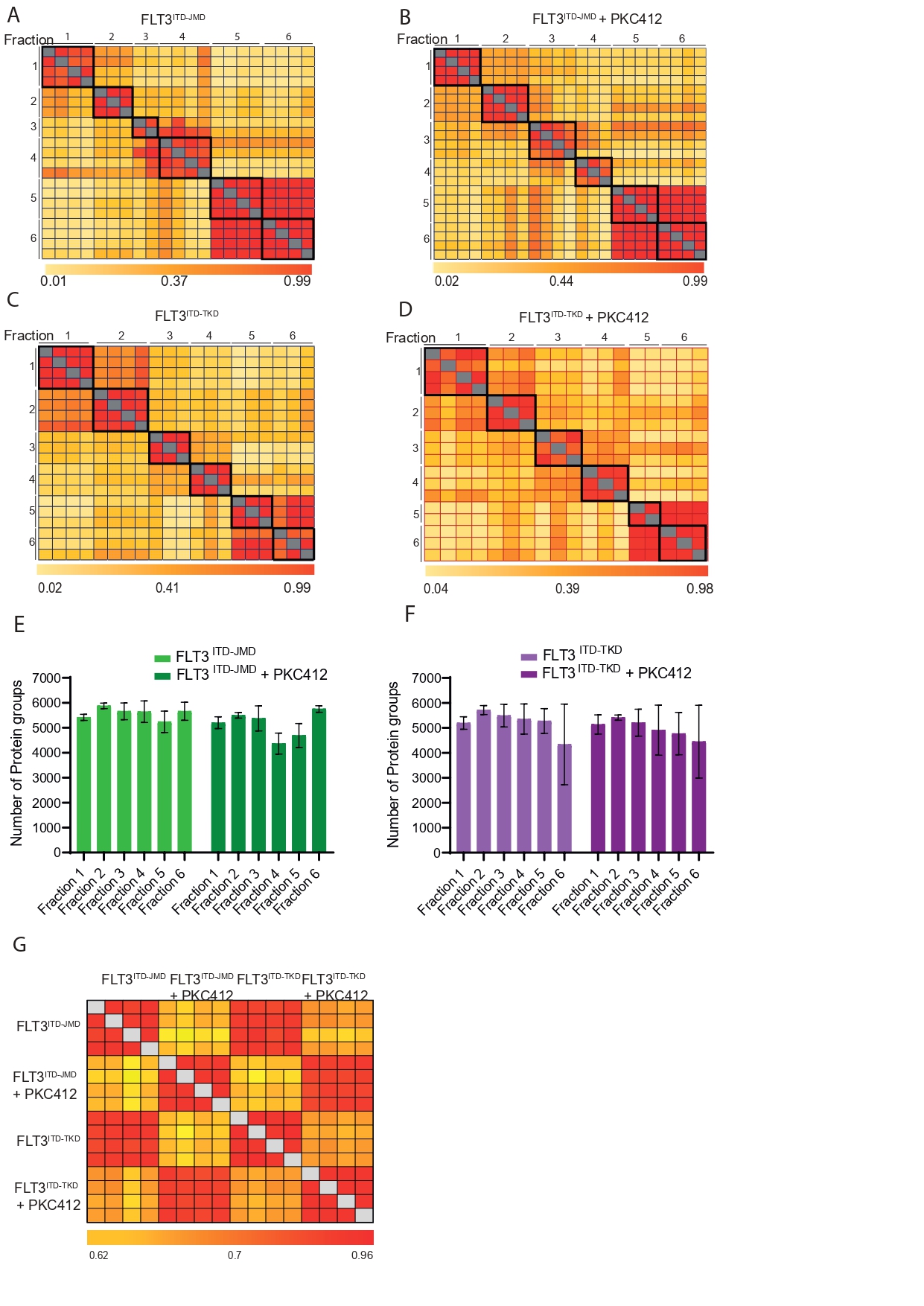

**Figure S1. (A-D)** Heatmaps reporting the Pearson Correlation coefficient of the proteomic analysis of the subcellular fractionation of control FLT3^ITD-JMD^(A), midostaurin-treated FLT3^ITD-JMD^ (B), FLT3^ITD-TKD^ (C) and midostaurin-treated FLT3^ITD-TKD^ (D). **(E-F)** Bar plots reporting the number of peptides identified in each biological condition and subcellular fraction in FLT3^ITD-JMD^ (E) and FLT3^ITD-TKD^ (F). **(G)** Heatmap reporting the Pearson Correlation coefficient of the phosphoproteomic analysis of control and midostaurin-treated FLT3^ITD-JMD^ and FLT3^ITD-TKD^.

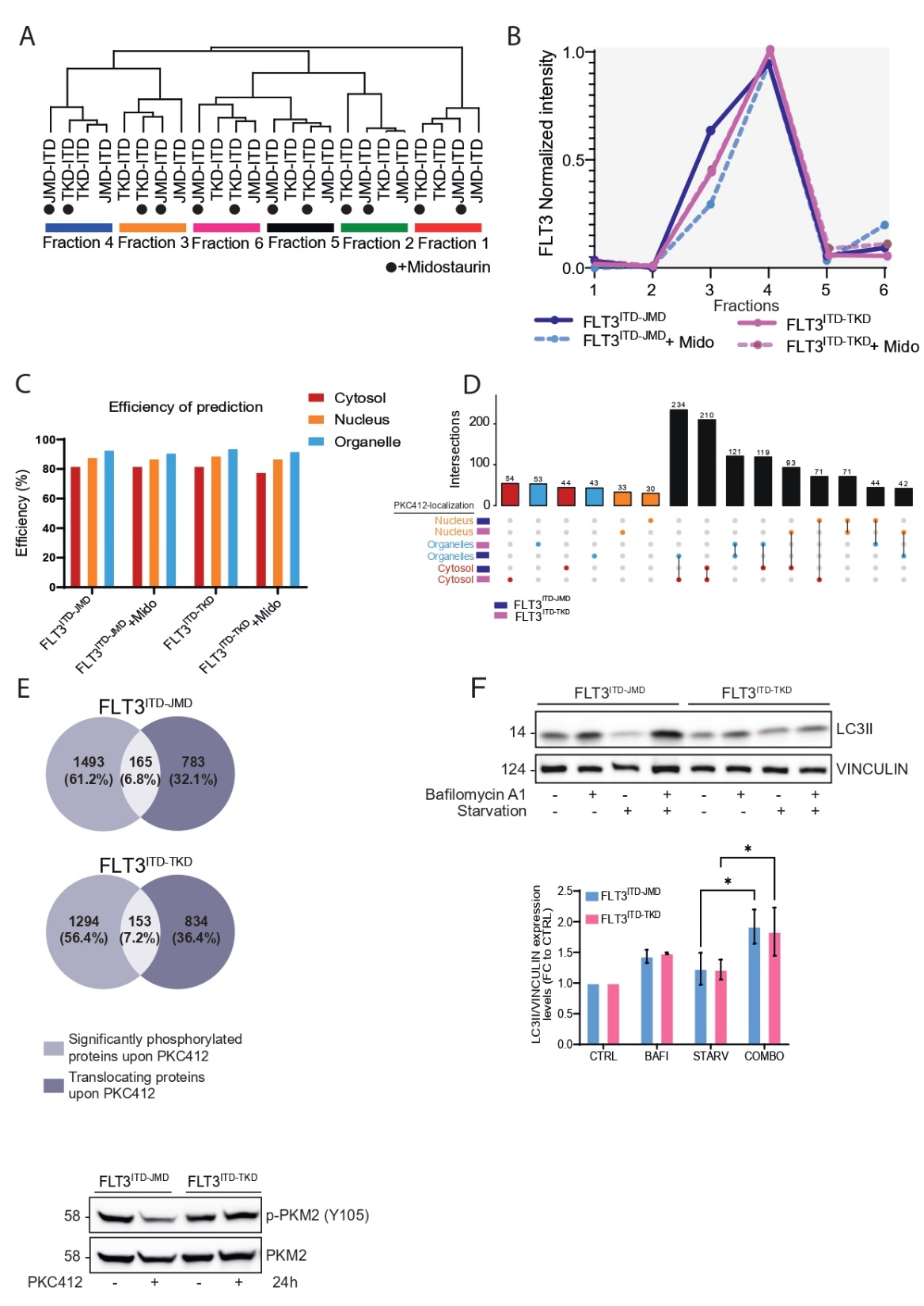

**Figure S2. (A)** Hierarchical clustering of subcellular fractionation data. **(B)** Intensity profiles of FLT3 protein in the six fractions (scaled intensities). **(C )** Bar plot reporting the percentage of the prediction efficiency of the SVM-organelle assignment. **(D)** Upset plot showing the number of translocating proteins that share the same subcellular location after midostaurin treatment in FLT3^ITD-JMD^ versus FLT3^ITD-TKD^ cells, or unique to each condition. **(E)** Venn diagrams reporting the percentage of translocating proteins whose phosphorylation is significantly modulated by midostaurin treatment or not. **(F)** Representative western blot and relative quantification of LC3-II protein levels after induction of autophagy by starvation in FLT3^ITD-JMD^ and FLT3^ITD-TKD^ either alone or combined with midostaurin treatment. (**G**) Representative western blot reporting the phosphorylation level of PKM2 at Y105 in FLT3^ITD-JMD^ and FLT3^ITD-TKD^ after 24 hours of 100nM midostaurin treatment.
