## Supplementary materials for "Organelle proteomics reveals novel metabolic vulnerabilities in FLT3-ITD cells"

**Immunoblot analysis**

BaF3 cells were seeded at a concentration of 500.000 cells/ml and treated as indicated. After treatments cells were centrifuged and washed in PBS 1x.  Next, cells were lysed in ice-cold lysis buffer (150 mM NaCl, 50 mM Tris–HCl, pH 7.5, 1% Nonidet P-40, 1 mM EGTA, 5 mM MgCl_2_, and 0.1% SDS) supplemented with 1 mM PMSF, 1 mM ortovanadate, 1 mM NaF, protease inhibitor mixture 1×, inhibitor phosphatase mixture II 1×, and inhibitor phosphatase mixture III 1× and incubated for 30 min. Protein lysates were separated at 13,000*g* for 30 min. The total protein concentration was determined using the Bradford reagent (Biorad, 5000006). Protein extracts were denatured and heated at 95°C for 10 min in NuPAGE LDS Sample Buffer (Thermo Fisher Scientfic, NP0007) and a buffer contained DTT as a reducing agent (NuPAGE Sample Reducing Agent) (Thermo Fisher Scientfic, NP0004). Proteins were resolved using 4–15% Bio-Rad Mini-PROTEAN TGX/CRITERION polyacrylamide gels (Bio-Rad 4561084). Proteins were transferred to Trans-Blot Turbo Mini Nitrocellulose Membranes using a Trans-Blot Turbo Transfer System (Bio-Rad, 17001918), and the nonspecific binding membranes were saturated in blocking solution (5% skimmed milk powder, 0.1% Tween 20 in 1× TBS) at room temperature for 1 hour. Saturated membranes were incubated overnight with primary antibodies diluted in BSA 5%. HRP-conjugated secondary antibodies (Goat Anti-Mouse/Anti-Rabbit IgG (H+L)-HRP Conjugate 1:3000, BIORAD 1721011) were diluted in blocking solution and used for the detection of the primary antibodies. Chemiluminescence was detected using Clarity Western ECL Blotting Substrates (Bio-Rad) and the Las-3000 Imaging System (Fujifilm). Band densities were quantified using ImageJ and normalized to the loading control.

**MTT assay**

Cell viability was measured using the Cell Proliferation Kit I (MTT) (Roche). Cells were treated as indicated. Then, MTT was added to the cells and incubated for 4 hours at 37 ◦C. Solubilization Solution was used to dissolve the formazan crystals during an overnight incubation. Finally, the plates were read at 590nm using a microplate reader (Bio-Rad).

**Immunofluorescence analysis**

Coverslips were incubated with poly-L-Lysine solution (Santa Cruz Biotechnology, sc-286689) for 1 hour at 37°C to allow cells’ adhesion. Upon treatment, cells were washed with 1x PBS and spotted on coverlips. Next, cells were fixed in 4% PFA for 15 min at RT and permeabilized with 0.3% Triton X-100 in 1x PBS for 10 min at RT. Unspecific bindings were saturated by incubating samples in blocking solution (3% BSA and 0.3% Triton X-100 in 1× PBS) for 1 hour. Cells were incubated with primary antibodies diluted in blocking solution according to manufacturer instruction. After incubation, cells were washed twice with 1x PBS and incubated, for 1h at RT, with host-specific secondary antibodies and 1:2000 DAPI solution (Thermo Scientific, 62248) diluted in blocking solution. Finally, the cells were washed twice with 1x PBS, mounted on slides and let dry overnight at RT.

**Subcellular fractionation**

Subcellular fractionation was performed following a recent in-house-developed spatial proteomics workflow (3) in biological quadriplicates. The workflow requires the preparation of three different lysis buffers:

| Buffer A | 30 mM Hepes pH 7.4; 15 mM NaCl, 2 mM MgCl2, 1 mM EDTA |
| --- | --- |
| Buffer B | 30 mM Hepes pH 7.4; 15 mM NaCl, 2 mM MgCl2, 1 mM EDTA, 350 mM sucrose |
| Buffer C | 30 mM Hepes pH 7.4; 15 mM NaCl, 2 mM MgCl2, 1 mM EDTA, 20% glycerol |

Protease and phosphatase inhibitors were added to each buffer at the final concentrations of: 1 mM PMSF, 1 mM ortovanadate, 1 mM NaF, protease inhibitor mixture 1×, inhibitor phosphatase mixture II 1×, and inhibitor phosphatase mixture III 1×.  n. Cell pellets were resuspended in 540 µl of Buffer B and 60 µl of 0.15% digitonin solution 5 min. Samples were incubated on orbital shaking at 4°C for 30 min and centrifuged for 3 min at 500 g. The recovered supernatant was marked as Fraction 1. Cell pellets were washed twice with 1 ml of buffer C. Cell pellets were resuspended in 540 µl of buffer B and 60 µl of 1.4 M NaCl. Samples were incubated on orbital shaking at 4°C for 30 min and centrifuged for 3 min at 500 g. The recovered supernatant was marked as Fraction 2. Cell pellets were washed twice with 1 ml of buffer B. Cell pellets were resuspended in 570 µl of buffer B and 30 µl of 10% Tween-20. Samples were incubated on orbital shaking at 4°C for 30 min and centrifuged for 3 min at 500 g. The recovered supernatant was marked as Fraction 3. Cell pellets were washed twice with 1 ml of buffer B. Cell pellets were resuspended in 540 µl of buffer C and 60 µl of 10% N-dodecyl maltoside. Samples were incubated on orbital shaking at 4°C for 30 min and centrifuged for 3 min at 500 g. The recovered supernatant was marked as Fraction 4. Cell pellets were washed twice with 500 µl of buffer C and resuspended in 540 µl of buffer A, 60 µl of 5 M NaCl and 1 µl of Benzonase® Nuclease. Samples were incubated on orbital shaking at 4°C for 30 min and centrifuged for 3 min at 500 g.  Supernatant was recovered and marked as Fraction 5. Cell pellets were washed once with 500 µl of buffer C and cell pellets were resuspended in 500 µl of buffer A 60 µl of 1.4 M NaCl and 18 µl of 10% SDS. Samples were immediately boiled for 10 min at 95 °C and marked as Fraction 6. Finally, all the fractions were centrifuged for 10 min at maximum speed.

**Samples preparation for proteomic analysis**

Protein precipitation in the 6 fractions was performed with methanol/chloroform precipitation and samples were resuspended in 2% SDC buffer in 100 mM Tris -HCl (pH 8.5). Proteins were reduced and alkylated with TCEP and CAA (1:100 v/v) at 45° for 5 minutes. Proteins digestion was performed adding trypsin and LysC enzymes (1:100 w/w) at 37° overnight. For the proteome preparation, we used the inStageTip (iST) method. Briefly, SDBRPS tips were washed with i) 100 µl acetonitrile (ACN), ii) 100 µl of 30% methanol and 1% TFA and iii) 150 µl of 0.2 % TFA centrifuging tips at 1000 xg for 3 minutes. Samples were loaded onto equilibrated columns and spin at 1000 xg for 10 minutes. SDBRPS tips were washed with i) 100 µl of 1% TFA, ii) 100 µl of 1% TFA in isopropanol and iii) 0.2% TFA. For the elution of proteins, we used a buffer containing 80% ACN, 5% NH_4_OH in MilliQ water. Samples were centrifuged at 1000 xg for 4 minutes and concentrated by SpeedVac at 45° for ~45 minutes. Finally, samples were dissolved in 10μl of a buffer containing 2% ACN and 0.1% TFA.

**Mass spectrometry analyses**

The peptides were desalted on StageTips and separated on a reverse phase column (50 cm, packed in-house with 1.9-mm C18- Reprosil-AQ Pur reversed-phase beads) (Dr Maisch GmbH) over 120 min or 140 min (single-run proteome and phosphoproteome analysis respectively). After elution, peptides were electrosprayed and analyzed by tandem mass spectrometry on a Orbitrap Exploris 480 (Thermo Fischer Scientific). The instrument was set to alternate between a full scan followed by multiple HCD based fragmentations scans for a total cycle time of up to 1 s.

**Proteome Data processing**

DIA Raw files were analyzed with Spectronauts HTRMS converter and analyzed with Spectronaut (v15.7.220308.50606). MS/MS spectra were matched against the *Mus musculus* UniProtKB FASTA database (September 2014), with an FDR of < 1% at the level of proteins, peptides and modifications. Enzyme specificity was set to trypsin, allowing for cleavage N-terminal to proline and between aspartic acid and proline. The search included cysteine carbamidomethylation as a fixed modification. Variable modifications were set to N-terminal protein acetylation and oxidation of methionine. Where possible, the identity of peptides present but not sequenced in a given run was obtained by transferring identifications across liquid chromatography (LC)-MS runs (‘match between runs’). Peptides had to be fully tryptic and up to two missed cleavages were allowed for protease digestion.

**Protein Correlation Profiling Analysis**

Proteomic analysis was performed in Perseus. LFQ intensities were scaled from 0-1. The intensity of proteins that were not quantified were set to 0. For normalization, the median values from the biological replicates for each fraction were calculated. Pearson correlations between the four biological replicates were calculated for each fraction. Annotations of proteins were extracted from UniProtKB and Gene Ontology (GO). Proteins with already known GO-cell compartment annotation and robust quantification were used as markers. A radial basis function (RBF) was used as the kernel. Protein markers were used to train the SVM based supervised learning approach implemented in Perseus. Sigma was set to 0.2 and C was set to 8. Organelle predictions were filtered for positive assignment to at least one organelle. Next, we estimated a second subcellular compartment contribution. We determined the highest correlation between the protein profile determined by our experiment with in silico generated combination profiles. The resulting correlation value is a proxy for the quality of secondary organelle assignment.

**Imaging cytometry**

Ba/F3 cells were cultured as described for 24 h in absence/presence of 100 nM PKC412 and harvested by centrifugation into tubes. Samples were stained with MitoViewTM 405 (biotium) according to manufacturer’s instructions. After nuclear counterstaining with 7AAD (Thermo Fisher) cells were analyzed on a Cytek FlowSight imaging cytometer and analyzed using IDEAS 6.2.

**Seahorse analysis**

Bioenergetics of Ba/F3 cells were analyzed after prior 24h culture in absence or presence of 100 nM PKC412. Cells were analyzed with the Seahorse XF Glycolysis Stress Test (Agilent), the Seahorse XF Mitochondrial Stress Test (Agilent) and the Seahorse XF Mito Fuel Flex Test (Agilent) according to the manufacturer’s recommendations on a Seahorse XFe 96 (Agilent) with 100.000 cells per well.

**Electron microscopy**

Ba/F3 cells expressing FLT3^ITD-JMD^ and FLT3^ITD-TKD^ constructs were fixed with 1% glutaraldehyde in 0.2 M HEPES buffer (pH 7.3) for 30 min at room temperature. Samples were washed three times with 1× PBS and centrifuged in the presence of 1% BSA for 10 min at 13,200 rpm until a compact cell pellet was obtained. Cell pellets were post-fixed for 30 min on ice with a mixture of 2% osmium tetroxide (OsO₄) and 3% potassium ferrocyanide. Subsequently, samples were incubated with 1% thiocarbohydrazide (TCH) diluted in H₂O for 5 min at room temperature. Pellets were then incubated overnight at 4 °C with 0.5% uranyl acetate diluted in ddH₂O. The following day, samples were dehydrated through a graded ethanol series (50%, 70%, 90%, and 100%) for 10 min at each step. Dehydrated pellets were infiltrated with a 1:1 mixture of acetone and EPON resin for 2 h, followed by incubation in pure EPON resin for an additional 2 h. Finally, samples were polymerized at 60 °C for 48 h. Three washes with ddH₂O were performed between each processing step. Serial ultramicrotomy was performed using a Leica EM UC7 ultramicrotome (Leica Microsystems), and thin sections (60 nm) were collected onto formvar carbon-coated slot grids. Imaging was performed using a Tecnai-12 transmission electron microscope (FEI) equipped with a VELETTA CCD digital camera (Soft Imaging Systems). Quantitative analysis of mitochondrial ultrastructure was performed using Fiji (ImageJ) image analysis software. Total mitochondrial area, the number of cristae per mitochondrion, and the cristae area normalized to the corresponding mitochondrial area were measured.
